## Supplementary File 2 for "Purine nucleosides replace cAMP in allosteric regulation of PKA in trypanosomatid pathogens"

Primers used in this study:

**pET-DUET TbPKAR(199-499) (expression of TbPKAR(199-499) in E. coli)**

TbPKAR199-499_FW	gccaGGATCCGgaaaacctgtattttcagGGATCTGGCCG GAAGCGACG
TbPKAR199-499_RV	cattatgcggccGCttaCTTCCTCCCCTCTGCCCT

**pET-DUET TbPKAR(199-499)_mutant 8 (expression of TbPKAR(199-499)_mutant 8 in E. coli)**

TbPKAR_M8_RV	atatatGCGGCCGCTTACTTCCTCCCCTCTGCCCTTAAAGTA

GTTTTCAGTTTCGACTGGGCGGCCTCATA

**pET-DUET TcPKAR(200-503) (expression of TcPKAR(200-503) in E. coli)**

TcPKAR200-503_FW	agccaGGATCCGgaaaacctgtattttcagGGATCTGGCAG AAACCGCCGCC
TcPKAR200-503_RV	cattatgcggccGCTTATACATCATCCACGTACGATG

**pET-DUET LdPKAR1(200-502) (expression of TcPKAR(200-503) in E. coli)**

LdPKAR1200-502_FW	TAGTGGATCCGGCCGGGCTCGTCGCCAGACAG

LdPKAR200-502_RV	attatgcggccGCCTACGCGCTCTCGTG

**Co-expression of TbPKAR-10xHis and Strep-TbPKAC1 in Lexsy (vectors pLEXSY_I-ble3, pLEXSY_I-neo3)**

5’PKAR SaII	ATGCATCTCGAGATGTCTGAAAAGGGAACATCG

3’PKAR NotI	TATATAATAGTTTAGCGGCCGCTCAGTGGTGGTGATGGTGGTGGT

GGTGGTGATGGCTACCGCCCTTCCTCCCCTCTGCCCTTAA

5’Strep BamHI	CGCGGATCCATGGCTTCGGCTTGGAGCCAC

PKAC1 Rev NotI	ACTATGCAATAGTTTAGCGGCCGCCTAAAAACCACGGAATGCAAC

5’Ble BamHI	GCCGGATCCACCATGGCCAAGTTGACCAGT

3’Ble SpeI	CTAGAACTAGTTCAGTCCTGCTCCTCGGCCA

**Co-expression of LdPKAR1-6xHis and Strep-LdPKAC1 in Lexsy (vectors pLEXSY_I-ble3, pLEXSY_I-neo3)**

LdPKAR_FW	ccgcctcgagatgggcagcagccatcaccatcatcaccacagccaggatccggaaaacctgtatttt

cagggatctATGTCCGCGGAAGACACCCCC

LdPKAR_RV	gagggcggccgccttaCTGGACCGCCGCCGGGGCACC

LdPKAC1_FW	gCCACCAgatctgCCATGGCTTCGGCTtggagccacccgcagttcgaaaaaGCTTCG

TCTGCTGCCAAGGACAGCTGTCCGGCGGA

LdPKAC1_RV	AGGAGGGCGGCCGCCTAAAAGCCATTGAATTCCGCCTGC

**Co-expression of TcPKAR-6xHis and Strep-TcPKAC2 in Lexsy (vectors pLEXSY_I-ble3, pLEXSY_I-neo3)**

TcPKAR_FW	ccgcctcgagatgggcagcagccatcaccatcatcaccacagccaggatccggaaaacctgtatttt

cagggatctATGTCCGCGGAAGACACCCCC

TcPKAR_RV	gagggcggccgccttaCTGGACCGCCGCCGGGGCACC

TcPKAC2_FW	gCCACCAgatctgCCATGGCTTCGGCTtggagccacccgcagttcgaaaaaGCTAC

GTTGAAGAATGCACCTGAATTTGTAAAGCCGGACGC

TcPKAC2_RV	AGGAGGGCGGCCGCTTAGAACCCAATAAACTCCGCCTGT

**Co-expression of TbPKAR-mutant1 and Strep-TbPKAC1 in Lexsy (vectors pLEXSY_I-ble3, pLEXSY_I-neo3)**

Mutant1_RBC-A_FW	GGAGAGCTTGcACTTATGTATCAGACACCAcgTGcTGCCACGGTG

Mutant1_RBC-A_RV	CACCGTGGCAgCAcgTGGTGTCTGATACATAAGTgCAAGCTCTCC

Mutant1_RBC-B_FW	GGTGAGCTGGcATTCCTTAACAATCACGCCcgTGcAGCAGATGTT

Mutant1_RBC-B_RV	AACATCTGCTgCAcgGGCGTGATTGTTAAGGAATgCCAGCTCACC

**Co-expression of TbPKAR-mutant2 and Strep-TbPKAC1 in Lexsy (vectors pLEXSY_I-ble3, pLEXSY_I-neo3)**

Mutant2_RBC-A_FW	GGAGAGCTCGcACTTATGTATCA

Mutant2_RBC-A_RV	TGATACATAAGTgCGAGCTCTCC

Mutant2_RBC-B_FW	AGCTGGcATTTTTAAACAAT

Mutant2_RBC-B_RV	ATTGTTTAAAAATgCCAGCT

**Co-expression of TbPKAR-mutant3 and Strep-TbPKAC1 in Lexsy (vectors pLEXSY_I-ble3, pLEXSY_I-neo3)**

Mutant3_RBC-A_FW	GTAGGAGAGCTTGAACTTATGTATCAGACACCggtTGTTGCCACG

Mutant3_RBC_A_RV	CACCGTGGCAACAaccGGTGTCTGATACATAAGTTCAAGCTCTCCTAC

Mutant3_RBC-B_FW	GTGGGTGAGCTGGAATTCCTTAACAATCACGCCgtTGTgGCAGAT

Mutant3_RBC-B_RV	ATCTGCcACAacGGCGTGATTGTTAAGGAATTCCAGCTCACCCAC

**Co-expression of TbPKAR-mutant4 and Strep-TbPKAC1 in Lexsy (vectors pLEXSY_I-ble3, pLEXSY_I-neo3)**

Mutant4_RBC-A_FW	CAGACACCAcgcGTCGCGACGGTG

Mutant4_RBC-A_RV	CACCGTCGCGACgcgTGGTGTCTG

Mutant4_RBC-B_FW	AATCACGCCcggGTAGCAGACGTCGTG

Mutant4_RBC-B_RV	CACGACGTCTGCTACccgGGCGTGATT

**Co-expression of TbPKAR-mutant5 and Strep-TbPKAC1 in Lexsy (vectors pLEXSY_I-ble3, pLEXSY_I-neo3)**

Mutant5_RBC-A-FW	GTAGGAGAGCTCGcACTTATGTATCAGACACCAcgcGTCGCGACGGTG

Mutant5_RBC-A-RV	CACCGTCGCGACgcgTGGTGTCTGATACATAAGTgCGAGCTCTCCTAC

Mutant5_RBC-B-FW	GGTGAGCTGGcATTTTTAAACAATCACGCCcggGTAGCAGACGTCGTG

Mutant5_RBC-B-RV	CACGACGTCTGCTACccgGGCGTGATTGTTTAAAAATgCCAGCTCACC

**Co-expression of human PKARIalpha and mouse PKACalpha (in pETDuet-1)**

5’strep (NdeI) Mm Cα	ttGGAATTCCATATGGCTTCGGCTtggagccacccgcagttcgaaaaaGCTGGCAAC

GCCGCCGCCGCCAAGAAGGGC

3’(AvrII)MmCα	ttcTGCCTAGGCTAAAACTCAGTAAACTCCTTGCC

6H.tev./HsPKARIaf	GCCAGGATCCGGAAAACCTGTATTTTCAGGGATCTATGGAGTCTGGCAGTA

6H.tev/HsPKARIaf	attatgcggccGCTCAGACAGACAGTGA
