## Supplementary material for "Purine nucleosides replace cAMP in allosteric regulation of PKA in trypanosomatid pathogens": Key resource table

**Key resources table**

| Reagent type (species) or resource | Designation | Source or reference | Identifiers | Additional information |
|---|---|---|---|---|
| Gene (*T. brucei*) | TbPKAR | TriTrypDB | Tb927.11.4610 | |
| Gene (*T. brucei*) | TbPKAC1 | TriTrypDB | Tb927.9.11100 | |
| Gene (*T. cruzi*) | TcPKAR1 | TriTrypDB | TcCLB.506227.150 | |
| Gene (*T. cruzi*) | TcPKAC2 | TriTrypDB | TcCLB.508461.280 | |
| Gene (*L. donovani*) | LdPKAR1 | TriTrypDB | LdBPK_130160.1 | |
| Gene (*L. donovani*) | LdPKAC1 | TriTrypDB | LINF_350045600 | |
| strain, strain background (*E. coli*) | Rosetta (DE3) | Novagen | SKU: 70954-3 | Electrocompetent cells |
| Cell line (*L. tarentolae*) | LEXSY T7-TR | Jena Bioscience | Cat.No.: LT-110 | |
| Cell line (*T. brucei*) | *T. brucei brucei* stock Lister 427 clone MiTat 1.2 | DOI: 10.1017/S0031182000046540 | | All T. brucei brucei cell lines are derived in the laboratory from T. brucei brucei stock Lister 427 clone MiTat 1.2, originally obtained from G. Cross, NY |
| Cell line (*T. brucei*) | *T.brucei* EATRO 1125 AnTat1.1 90:13 | DOI: 10.1101/gad.323404 | | |
| Cell line (*T. brucei*) | MITat1.2SM 6HIS TbPKAR | This paper | | MITat1.2_SM blood stream forms (BSF) of *T. brucei* with expression of TbPKAR fused to a 6xHis tag |
| Cell line (*T. brucei*) | EATRO11252T7 6HIS TbPKAR | This paper | | EATRO11252T7 procyclic forms (PCF) of *T. brucei* with expression of TbPKAR fused to a 6xHis tag |

| | | | | |
|---|---|---|---|---|
| Cell line (*T. brucei*) | MITat1.2SM PKAR-KO | DOI: 10.1038/s41467-019-09338-z | | MITat1.2_SM BSF PKAR knock out |
| Transfected construct (*T.brucei*) | pLEW100v5b1d-BSD_6His-Tev TbPKAR | This paper | | Construct cloned and transfected into MITat1.2SM (BSF) |
| Transfected construct (*T.brucei*) | pHD1146-puro_6xHis-Tev TbPKAR | This paper | | Construct cloned and transfected into EATRO11252T7 (PCF) |
| antibody | anti-PKAR (rabbit) | DOI: 10.1038/s41467-019-09338-z | | 1:500 |
| antibody | Anti-PKAC1/2 (rabbit) | DOI: 10.1038/s41467-019-09338-z | | 1:500-1:1000 |
| antibody | Anti-6x His tag (mouse) | Thermofisher scientific | | 1:1000 |
| antibody | IRDye® 800CW anti-mouse (goat) | LICOR | Cat# 925-32210 | 1:5000 |
| antibody | IRDye® 680LT anti-rabbit (goat) | LICOR | Cat# 925-69021 | 1:5000 |
| antibody | Alexa Fluor® 680 anti-rabbit (goat) | ThermoFisher | Cat# A27042 | 1:5000 |
| Recombinant DNA reagent | pLEXSY_I-ble3 (vector) | Jena Bioscience | Cat# EGE-244 | |
| Recombinant DNA reagent | pLEXY_I-neo3 (vector) | Jena Bioscience | Cat# EGE-245 | |
| Recombinant DNA reagent | pLEXSY_I-ble3_TbPKAR (plasmid) | DOI: 10.1038/s41467-019-09338-z | | Transfected in LEXSY expression system |
| Recombinant DNA reagent | pLEXSY_I-ble3_TbPKAR_mutant 1-7 (plasmids) | This Paper | | Transfected in LEXSY expression system; for specific mutation inserted see Table 2 |
| Recombinant DNA reagent | pLEXSY_I-ble3_TcPKAR1 (plasmid) | This paper | | Transfected in LEXSY expression system |
| Recombinant DNA reagent | pLEXSY_I-ble3_LdPKAR1 (plasmid) | This paper | | Transfected in LEXSY expression system |

| | | | | |
|---|---|---|---|---|
| Recombinant DNA reagent | pLEXY_I-neo3_TbPKAC (plasmid) | DOI: 10.1038/s41467-019-09338-z | | Transfected in LEXSY expression system |
| Recombinant DNA reagent | pLEXY_I-neo3_TcPKAC2 (plasmid) | This paper | | Transfected in LEXSY expression system |
| Recombinant DNA reagent | pLEXY_I-neo3_LdPKAC1 (plasmid) | This paper | | Transfected in LEXSY expression system |
| Recombinant DNA reagent | pETDuet-1 DNA-Novagen (vector) | Sigma-Aldrich Novagen | SKU 71146 | |
| Recombinant DNA reagent | pET_SUMO Expression system | ThermoFisher Scientific | K30001 | |
| Recombinant DNA reagent | pETDuet-1_TbPKAR(199-499) | DOI: 10.1038/s41467-019-09338-z | | Recombinant expression of TbPKAR(199-499) in *E. coli* |
| Recombinant DNA reagent | pETDuet-1_TbPKAR(199-499)_mutant 6 | This paper | | Recombinant expression of TbPKAR(199-499) mutant 6 in *E. coli* |
| Recombinant DNA reagent | pETDuet-1_TbPKAR(199-499)_mutant 7 | This paper | | Recombinant expression of TbPKAR(199-499) in *E. coli* |
| Recombinant DNA reagent | pETDuet-1_TbPKAR(199-499)_mutant 8 | This paper | | Recombinant expression of TbPKAR(199-499) in *E. coli* |
| Recombinant DNA reagent | pETDuet-1_TcPKAR1(200-503) | DOI: 10.1038/s41467-019-09338-z | | Recombinant expression of TcPKAR(200-5003) in *E. coli* |
| Recombinant DNA reagent | pETDuet-1_HsPKAR1$\alpha$ | PMID: 8393867 | | Recombinant expression of HsPKAR1$\alpha$ in *E. coli* |
| Recombinant DNA reagent | pET-11_Sumo3_LdPKAR1(200-502) | This paper | | Recombinant expression of LdPKAR1(200-502) in *E. coli* |
| Peptide, recombinant protein | TEV Protease | NEB | Cat# 8112S | Cleavage of HIS-Tag in recombinant protein |
| Peptide, recombinant protein | Sumo Protease | Sigma-Aldrich | SKUSAE 0067-2500UN | Cleavage of SUMO Tag in recombinant protein |
| Commercial assay or kit | Hi Yield® Plasmid Mini DNA Isolationkit | Süd-Laborbedarf GmBH, Germany | Art-Nr.: 30 | |

| | | | |
|---|---|---|---|
| | | | HYPD100 |
| Chemical compound, drug | Inosine | Sigma Aldrich | Cat# 200-390-4 |
| Chemical compound, drug | Guanosine | Sigma Aldrich | Cat# 204-227-8 |
| Chemical compound, drug | Adenosine | Sigma Aldrich | Cat# 93029 |
| Chemical compound, drug | cAMP | Biolog Life Science Institute | Cat# A 001 H |
| Chemical compound, drug | cGMP | Biolog Life Science Institute | Cat# G 001 |
| Chemical compound, drug | cIMP | Biolog Life Science Institute | Cat# I 001 |
| Chemical compound, drug | AMP | Sigma Aldrich | Cat# 54612 |
| Chemical compound, drug | GMP | Santa Cruz Biotechnology | Cat# 226-914-1 |
| Chemical compound, drug | IMP | Sigma Aldrich | Cat# 57510 |
| Chemical compound, drug | 2'-deoxyadenosine | Sigma Aldrich | Cat# D7400-250MG |
| Chemical compound, drug | 3'-deoxyadenosine | Sigma Aldrich | Cat# C3394 |
| Chemical compound, drug | 5'-deoxyadenosine | Sigma Aldrich | D1771 |
| Chemical compound, drug | Nebularine | Santa Cruz Biotechnology | Cat# sc-208087 |
| Chemical compound, drug | Allopurinol riboside | Santa Cruz Biotechnology | Cat# sc-217610 |
| Chemical compound, drug | Xanthosine | Sigma Aldrich | Cat# X0750 |
| Chemical compound, drug | Uridin | Sigma Aldrich | Cat# U3750 |

| | | | | |
|---|---|---|---|---|
| Chemical compound, drug | Cytidin | Sigma Aldrich | Cat# C122106 | |
| Chemical compound, drug | 13C5-labeled Inosin | Omicron Biochemicals Inc | | Doi 10.1038/s41596-018-0094-6 |
| Chemical compound, drug | 13C5-labeled Guanosin | Omicron Biochemicals Inc | | Doi 10.1038/s41596-018-0094-6 |
| Chemical compound, drug | 13C5-labeled Adenosin | Omicron Biochemicals Inc | | Doi 10.1038/s41596-018-0094-6 |
| Software, algorithm | GraphPad Prism 7.0 | GraphPad | | Statistical testing |
| Software, algorithm | Phenix | doi:10.1107/S0907444909052925 | | Model refinement |
| Software, algorithm | Coot | doi:10.1107/S0907444910007493 | | Manual model building of protein structure |
| Software, algorithm | Glide (Maestro) | Schroedinger LLC, New York, NY, 2023 | | Molecular docking |
| Software, algorithm | The PyMOL Molecular Graphics System (Version 2.0) | Schrödinger, LLC | | Illustration of structural figures |
